# Medial entorhinal cortex activates in a traveling wave

**DOI:** 10.1101/632109

**Authors:** J.J. Hernández-Pérez, K.W. Cooper, E.L. Newman

## Abstract

Traveling waves of cortical activity are hypothesized to organize cortical information processing and support interregional communication. Yet, it remains unknown whether interacting areas exhibit the matched traveling waves necessary to support this hypothesized form of interaction. Here, we show that the strongly-interacting medial entorhinal cortex (MEC) and hippocampus exhibit matched traveling waves. We demonstrate that both the field potential and spiking in the MEC exhibit prominent 6-12 Hz ‘theta’ traveling waves matching those of the hippocampus. The theta phase shifts observed along the MEC were accounted for largely by variation in waveform asymmetry. From this, we hypothesize that that gradients in local physiology underlie both the generation of MEC traveling waves and the functional variations observed previously across the MEC.

## Introduction

A major outstanding challenge in modern neuroscience is to bridge from physiology to function. Extracellular field potential recordings scaffold this bridge, enabling linkage between biomarkers of functional processes (e.g., memory encoding or retrieval) and the underlying physiology. Yet, limiting this bridge is a lack of knowledge regarding the broad spatiotemporal dynamics that organize activity across the relevant circuits. What is needed now is to understand how those biomarkers manifest at the *whole-circuit* scale.

The recent recognition that numerous cortical circuits exhibit traveling waves when engaged motivates the hypothesis that traveling waves are a basic organizing principle of cortical activity (Muller et al., 2018). Traveling waves, by this hypothesis, facilitate interaction between functionally-distinct circuits by generating macroscopic structure in precise activation dynamics across anatomically distributed circuits. Critically, the functionality of traveling waves is distinct from the underlying mechanism. They need not be generated by a specific mechanism (e.g., locally propagating activity) to generate macroscopic structured activity, rather, they must exhibit matched traveling wave dynamics. To date, however, traveling waves have only been observed in isolated circuits (c.f., van Ede et al., 2015). A goal of this work was to test if matched traveling waves exist in two functionally-distinct strongly-interacting circuits, the hippocampus and the entorhinal cortex.

The hippocampus exhibits traveling waves that are understood to be critical for the functionality of the circuit. The ∼8 Hz ‘theta’ rhythm that is readily observed at individual electrodes travels from the dorsal/septal pole to the ventral/temporal pole of the hippocampus in humans and rats alike (Lubenov and Siapas, 2009; Patel et al., 2012; Zhang and Jacobs, 2015). Theta is well understood both in terms of its underlying physiology and its potential function (for reviews, see Buzsáki, 2002; Dickson et al., 2000; Vinogradova, 1995). Functionally, theta is attributed with supporting navigation (Blair et al., 2007; Brandon et al., 2011; Burgess et al., 2007; Hasselmo et al., 2009; Newman and Hasselmo, 2014; Onslow et al., 2014), mediating associational learning (Caplan et al., 2003; Hasselmo et al., 2002; Hernández-Pérez et al., 2015; 2016; Honey et al., 2017; Muller et al., 2018; Norman et al., 2006; Patel et al., 2012), and pacing the relative timing of the distinct components of the entorhinal-hippocampal circuit (Mizuseki et al., 2009). These proposed functions are not mutually exclusive. However, for each, hippocampal input from the entorhinal cortex must also be a traveling wave.

The medial entorhinal cortex (MEC), like the hippocampus, exhibits clear extracellular field theta rhythms (Mitchell and Ranck, 1980). Simultaneous recordings of theta in the MEC and hippocampus demonstrate a stable phase offset between these areas in terms of theta phase (Mitchell and Ranck, 1980; Mizuseki et al., 2009) consistent with the idea that theta facilitates interaction between these areas. At individual points along the dorsal-ventral axis of the MEC, theta in layers III-V is largely synchronous but reverses phase between layers III and I (Chrobak and Buzsáki, 1998; Mitchell and Ranck, 1980; Quilichini et al., 2010). At the point of inflection in layer II, theta power drops to zero. It is unknown, however, whether theta phase varies systematically along the dorsal-ventral axis of the MEC.

Here, we aimed to establish whether matched traveling waves exist in the hippocampus and the functionally adjacent MEC. To do so, we used custom electrode arrays to record the field potential at regular intervals along the dorsal-ventral axis of the MEC in freely behaving rats. Our recordings revealed reliable phase shifts across the dorsal-ventral extent of the MEC indicating the existence of traveling waves. We further analyzed these recordings for evidence regarding the mechanism by which these traveling waves were generated. These analyses revealed that the waveform of theta shifted gradually across the dorsal-ventral axis, accounting for much of the observed phase shifts indicating that theta traveling waves are generated by varying local physiology.

## Results

### Theta phase shifts gradually across the dorsal-ventral axis of the MEC

To determinate whether there is a traveling wave in the MEC, we used custom-built electrode arrays to recorded extracellular field potentials at regular intervals along the dorsoventral axis of MEC while the rats completed laps on a circle track for food rewards (Figure 1). In each of the six rats implanted, gradual phase shifts in the 6-12 Hz theta-band were apparent in the raw recordings. In each, dorsal sites led relatively more ventral sites (e.g., Figure 1C). This is illustrated by computing cycle triggered averages of the raw LFP triggering on theta peaks at the dorsal-most electrode (Figure 2A).

**Figure 1.**
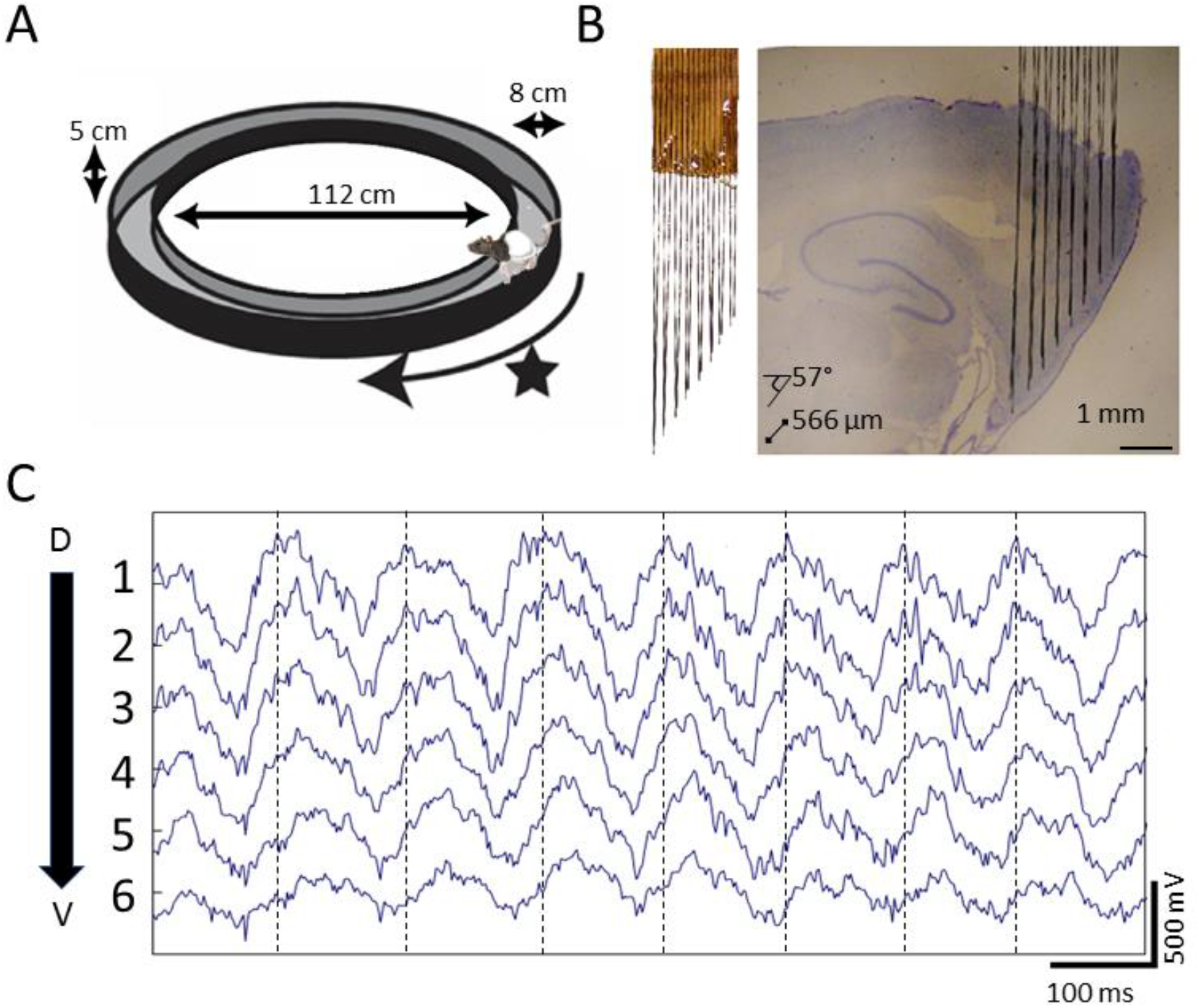
Extracellular field potential recordings along ∼3mm of the dorsal-ventral axis of the MEC in behaving rats reveals a gradual phase shift of theta. A) Rats completed laps on a circle track for food rewards delivered at a fixed location marked by the star. B) Custom electrode arrays with fixed interelectrode spacing (566 μm and 57° orientation relative to horizontal plane) were used to sample the field potential along ∼3 mm of the long axis of the MEC. Array depth was controlled by a micro-drive (not shown). C) Example broadband LFP traces from adjacent electrodes in MEC reveal regular phase shifts. Vertical lines mark the theta peaks from the dorsal-most channel to facilitate visual comparison of phase shifts across channels.

**Figure 2.**
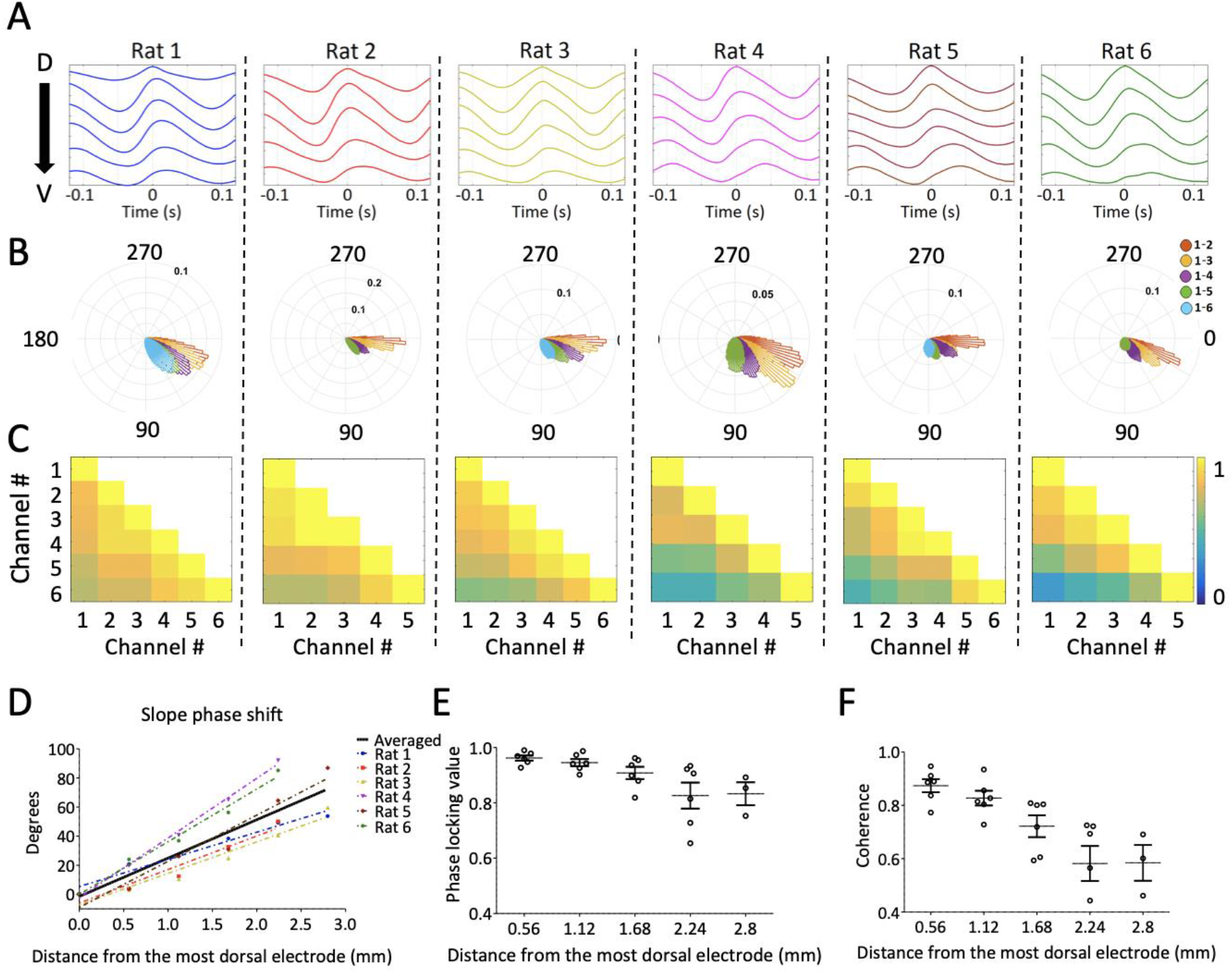
Consistent gradual phase shifts were observed across along the long axis of MEC in all animals. A) Average theta waves across channels, computed as an event related average of the raw LFP triggered on the peak of theta on the dorsal-most channel, show similar theta phase shifts across channels for each rat. B) Histogram of theta phase differences for each electrode relative to the dorsal-most electrode for each rat. C) Theta-band coherence between all electrode-pairs for each rat. Electrode one corresponds to the most dorsal position. D) Summary of theta phase shifts of each electrode relative to the dorsal-most electrode as a function of the distance between the electrodes. See also Figure S2. E) Phase locking (mean resultant length) between each electrode and the dorsal-most electrode, plotted as a function of inter-electrode distance, shows consistently high phase-locking across the dorsal-ventral axis. F) Summary of theta-band coherence changes between each electrode and the dorsal-most electrode (corresponding to left-most column of matrices shown in C) plotted as a function of inter-electrode distance. Error bars in E and F reflect standard errors.

Theta-phase changed linearly as a function of distance between the recording sites (Figure 2A, 2D). The phase offset between any two channels was relatively stable over the duration of the of the recording indicating strong phase locking between sites. Histograms of the phase differences are shown in Figure 2B. Plotting phase locking as a function of distance between electrodes (Figure 2E) revealed only a modest decrease at the maximum separation with most pairs exhibiting phase locking (in terms of mean resultant length) of greater than 0.8. Consistent with this, theta coherence was also generally high across the MEC but decreasing as the distance between electrode pairs increased (Figure 2C, 2F). These patterns are consistent with previously reported shifts in theta phase and coherence across the long axis of the hippocampus (Lubenov and Siapas, 2009; Patel et al., 2012) suggesting similar traveling waves in both structures. From these observations, we conclude that there are theta-band traveling waves in the MEC.

Next, we sought to determine the total theta phase shift across the dorsal-ventral axis of the MEC. With our recordings, we observed a shift of ∼65° across the dorsal-most ∼2.5 mm of the MEC. This corresponds to an average shift of ∼26 °/mm. The total length of the MEC in adult rats is 6-7 mm (Insausti et al., 1997; Long et al., 2015). Assuming the phase shift per millimeter is consistent across the remaining length of the MEC, the total theta phase shifts would be 155-180°. In the hippocampus, the phase of theta shifts by 180° between the dorsal and ventral poles (Patel et al., 2012). This would indicate that the MEC and hippocampus have similar overall phase shifts along their dorsal-ventral axes.

To rule out the possibility that the observed phase shifts could be attributed to variable placement of the electrode tips with respect to the cortical layers of MEC, we implanted one rat with a variant of the electrode array that additionally sampled across the layers of the MEC (Figure S2). This enabled direct comparison of theta phase differences across versus along the layers of MEC. Because the positions of individual electrodes were fixed relative to other electrodes in the array, we could determine the displacement for each electrode-pair with respect to an axis running perpendicular (i.e., spanning) the layers and with respect to an axis running parallel (i.e., along) the layers. Then for each electrode pair, we plotted the observed theta phase shift as a function of the displacement along one axis or the other (Figure S2 C-F). This revealed a clear positive correlation between phase-offset and with distance ‘along the cell layer’ (rho = 0.58). In contrast, there was no correlation between phase-offset and distance ‘across the layers’ (rho = 0.02). From this, we conclude that the reported phase shifts cannot be attributed to inconsistent placements

Next, we sought to verify that the phase shifts observed in the field potential were indicative of the local neural activity. To do so, we shifted the electrodes into layer II where there is strong multi-unit activity and asked whether the timing of multi-unit activity mirrored that of the field potential. In three of the animals, clear multi-unit activity was observable simultaneously across four or more of the channels when the probe was lowered into layer II. As expected from prior work, this multiunit activity exhibited clear theta-rhythmicity (Mitchell and Ranck, 1980). The amplitude envelope of the 600-3000 Hz spiking-band activity showed a progressive phase shift across channels resembling that which we observed in the theta field potential (Figure 3). Computing the cross-correlogram of the amplitude envelope between each channel and the dorsal-most channel revealed an increasing lag matching the lags observed in the field potential (Figure 3D). These results demonstrate that neural activity, like the field theta, exhibited traveling wave dynamics along the dorsal-ventral axis.

**Figure 3.**
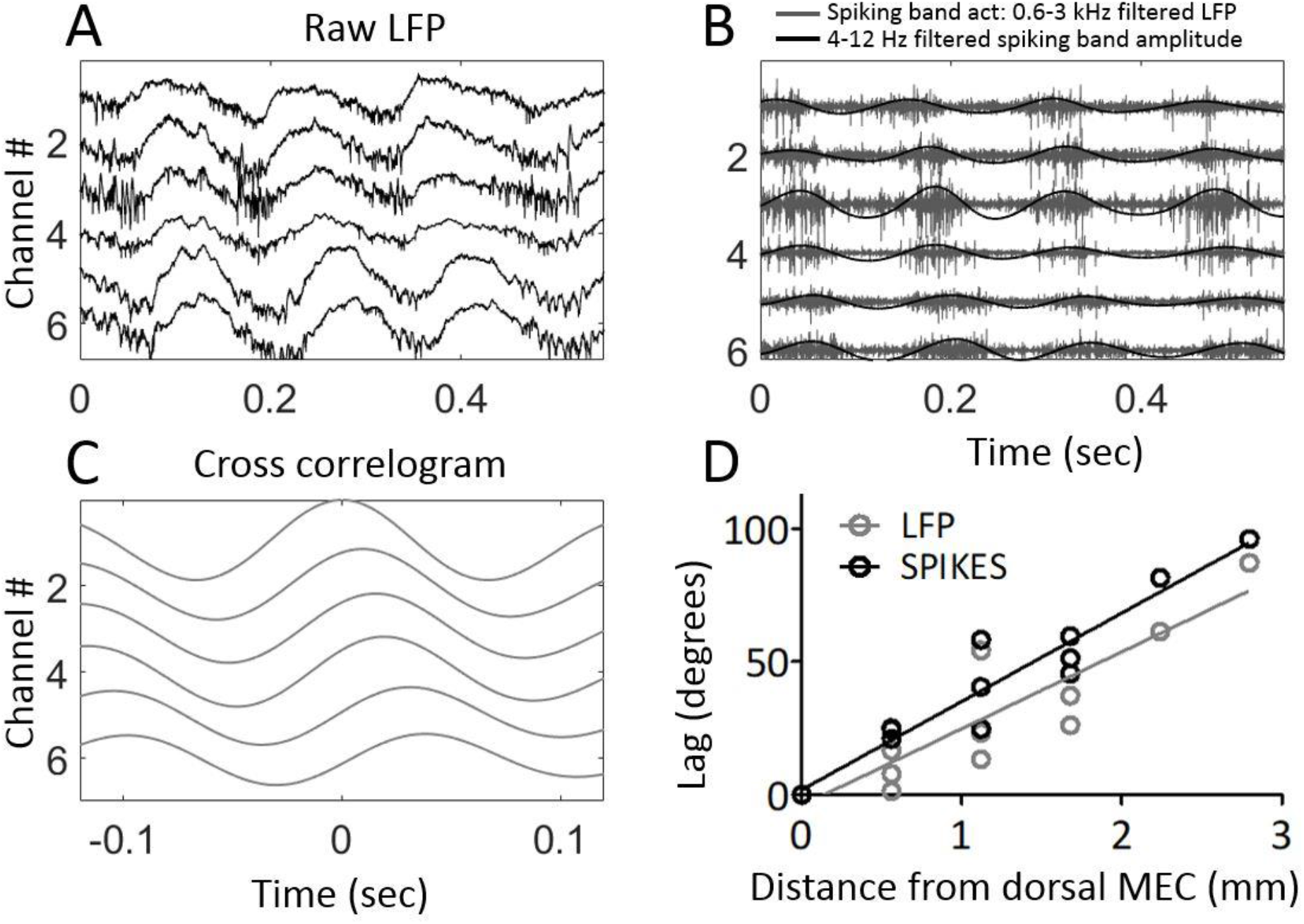
Multi-unit activity exhibited traveling waves along the dorsal-ventral axis. A) Representative example of multi-unit activity locked to the trough of the local theta that was observable in the broadband LFP when the electrode arrays were lowered into MEC layer II. B) Bandpass filtering the broadband signal to 600-3000 Hz to focus on the ‘spiking-band,’ shown in gray, shows that the timing of these theta-rhythmic fluctuations in multi-unit activity varied across the dorsal-ventral axis of the MEC in this representative example. Low-pass filtering of the spiking-band amplitude, shown in black, highlights the theta rhythmic structure of the spiking activity. C) Cross-correlations of the spiking-band amplitude between the dorsal-most channel and each of the other channels from the same trial as shown in A and B resemble the traveling LFP theta waves. D) A consistent pattern of increasing lags was observed across the three rats with multiunit activity on four or more channels simultaneously. The lag to the cross-correlation peaks (black points and best-fit line) increases as a function of distance from the dorsal-most electrode. The slope closely mirrors the slope in LFP theta phase offsets (grey points and best-fit line). The linearly increasing lags confirm that multi-unit activity activates at progressively longer delays along the dorsal-ventral axis.

### Theta waveform changes are a source of phase shift along the dorsal-ventral axis of MEC

Visual inspection of theta revealed a clear asymmetric waveform in dorsal MEC (e.g., Figure 1C). The strength of this asymmetry varied along the dorsal-ventral axis. It was strongest at dorsal sites and faded at ventral recording sites (Figures 4). To analyze these changes, we compared the duration of the rising phase to the duration of the decaying (falling) phase and the duration of the peak to the duration of the trough using a variant of the waveform asymmetry index (Belluscio et al., 2012; Cole and Voytek, 2017).

**Figure 4.**
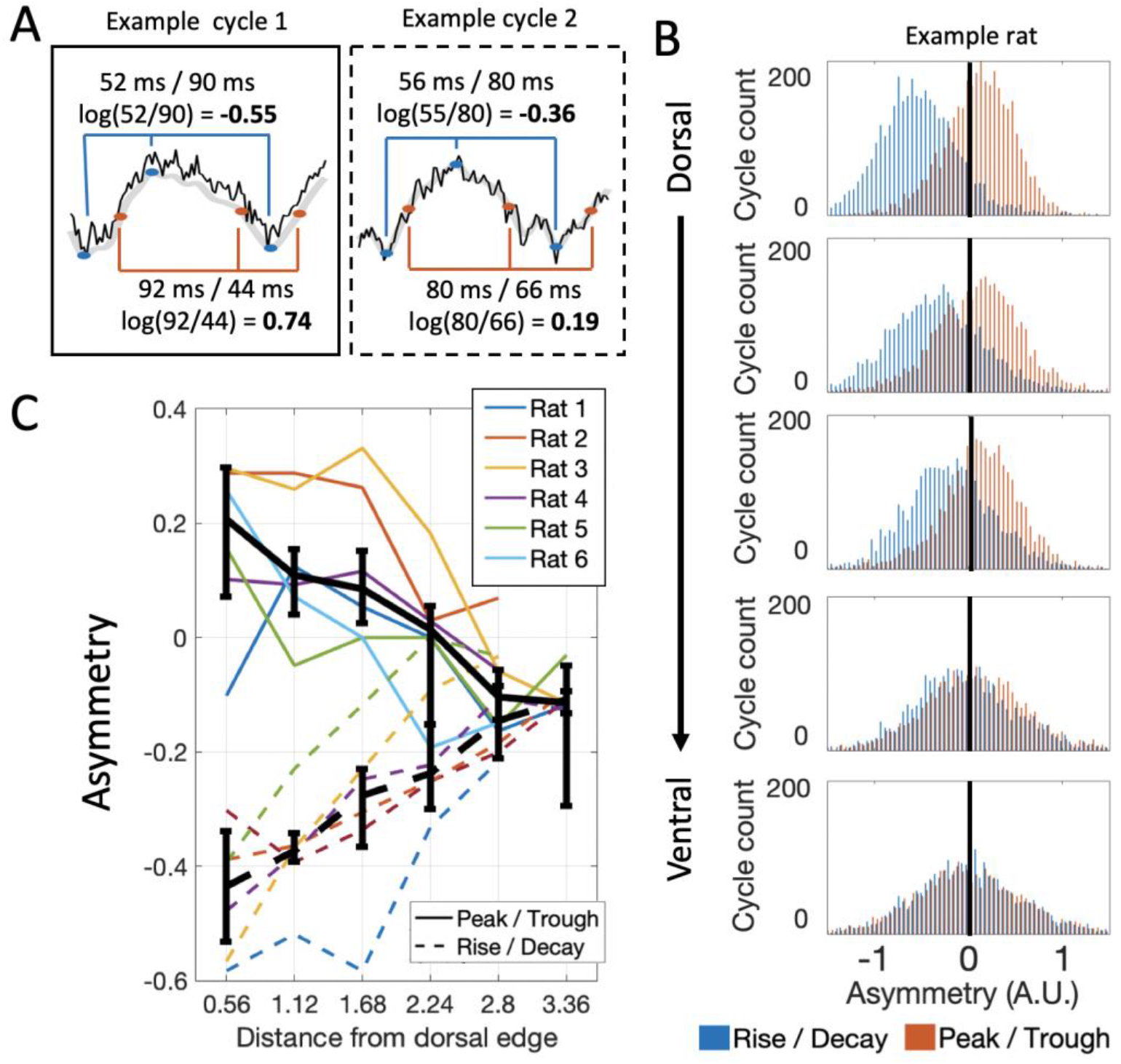
Theta waveform asymmetry varies along the dorsal-ventral axis of the MEC. A) How waveform asymmetry was analyzed. Top – the rise/decay asymmetry was calculated by measuring the duration of the rising and decaying (falling) phases (*e.g.1, 52 and 90 ms*, respectively; marked with blue lines) and taking the log of the ratio of these durations (*e.g.1, log(52/90) = −0.55*). Bottom - the peak/trough asymmetry was calculated likewise: as the log of the ratio of the peak and trough durations (*e.g.1, log(92/44) = 0.74*; marked with red lines). B) Representative example of rise/decay asymmetry and peak/trough asymmetry distributions observed over all theta cycles recorded across channels in a single trial from Rat 4. The rise/decay asymmetry (blue bars) at dorsal sites was dominantly negative indicating a fast rise and slow decay sawtooth waveform. Ventral sites were less negative indicating more symmetric rise and decay times. The peak/trough asymmetry (orange bars) at dorsal sites was mostly positive indicating that peaks were longer than troughs. Ventral sites were less positive indicating more symmetric peak and trough durations. C) Summary over rats. Colored lines show the median asymmetry values for individual rats. Black lines show the median across rats with 95% bootstrap confidence intervals on the within subject effect of electrode position. The rise/decay asymmetries (dashed lines) and peak/trough asymmetries (solid lines) start at respective extremes at dorsal sites and converge on a slight negative value ventrally. This indicates that dorsal theta has a prominent sawtooth pattern that diminishes ventrally and that the troughs grow progressively longer at ventral sites relative to dorsal sites.

In dorsal MEC, theta was characterized by a rapid rise and slow decay as indicated by reliably negative rise/decay asymmetry values (median [95% CI] = −0.43 [-0.57 −0.35], signed rank [max] = 0 [21], p = 0.03, n = 6; Figure 4C). This asymmetry decreased significantly across the dorsal-ventral axis (rank sum [max] = 0 [21], p = 0.03, n = 6; Figure 4C). Yet, even at the ventral sites, the rise/decay asymmetry values remained reliably negative (at ∼2.80 mm from dorsal edge of MEC: median [95% CI] = −0.11 [-0.13 −0.09]; signed rank [max] = 20 [21], p = 0.06, n = 6).

Asymmetries in peak and trough duration also varied across the dorsal-ventral axis. Theta in dorsal MEC was characterized by relatively long peaks and short troughs. In five of six animals, the peak/trough asymmetry values were positive (median [95% CI] = 0.21 [0.00 0.29]; signed rank [max] = 20 [21], p = 0.06, n = 6; Figure 4C;). In all animals, the peak/trough asymmetry shifted toward negative values between the dorsal and ventral portions of MEC (rank sum [max] = 0 [21], p = 0.03, n = 6; Figure 4). Though not significant in this sample size, the peak/trough asymmetry values showed a trend toward negative values at ventral sites (at 2.80 mm from dorsal edge of MEC: median [95% CI] = −0.10 [-0.16 0.01]; signed rank [max] = 3 [21], p = 0.16, n = 6).

A consequence of varying waveform is that the time required for a theta wave to travel from dorsal sites to ventral sites (i.e., conduction delay) varies across phases. To characterize this, we computed the conduction delay that results when we restrict our analysis to specific reference phases: the falling phase (mid-point between the peak and trough), the trough, the rising phase (mid-point between trough and peak), and the peak (90°, 180°, 270°, 360° respectively). We found that the conduction delay was shortest for the falling phase (median [95% CI] = 5.47 ms/mm [2.87 8.70]; Figure 5B) and greatest for the peak to travel from dorsal sites to ventral sites (median [95% CI] = 12.00 ms/mm [6.52 14.43]; Figure 5B). Across rats, the conduction delay at the falling phase was reliably shorter than at the peak (rank sum [max] = 0 [21], p = 0.03, n = 6), with about half of the delay (5.47 vs. 12.00 ms/mm). The conduction delay increased steadily between the falling phase and the peak (Figure 5C). Fitting a line to the delays observed across phases spanning 90 to 360 revealed a significantly positive slope of 0.02 (ms/mm)/deg (95% C.I. = [0.017 0.025]; R^2^ = 0.82; Figure 5C). These results show that theta is most synchronized during the falling phase and becomes progressively more desynchronized across phases until the peak of theta.

**Figure 5.**
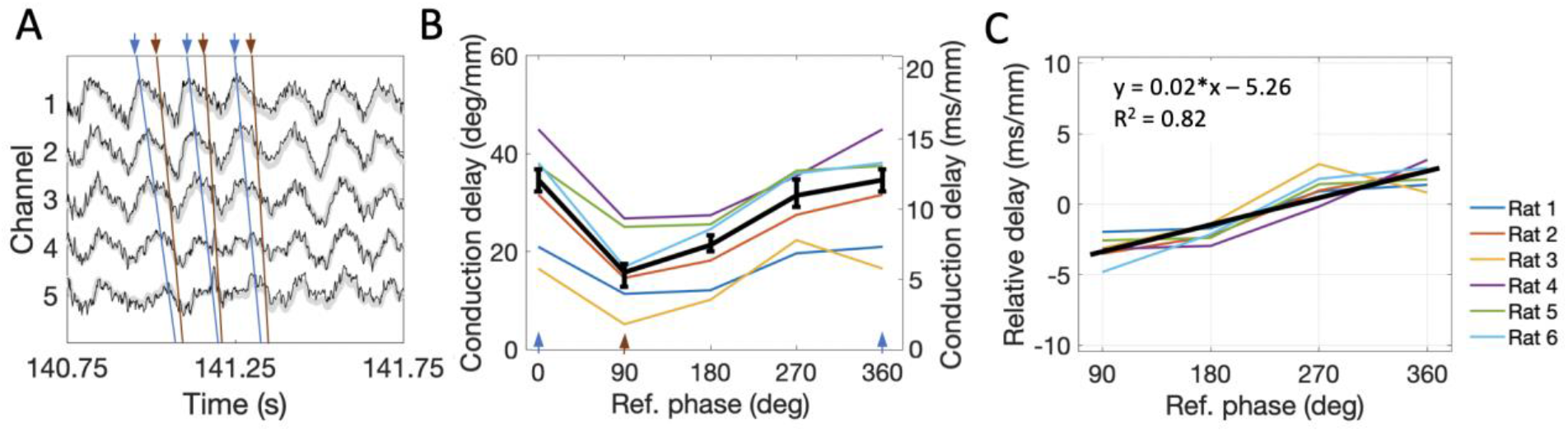
The conduction delay for theta to travel along the dorsal-ventral axis depends upon what phase is used as a reference point. A) Representative snip of LFP. Blue lines connect the peaks across channels and orange lines connect falling phases across channels. The orange lines appear steeper than the blue lines indicating shorter conduction delays. B) Across rats, the conduction delay is longest for theta peaks (0 or 360 deg, marked with blue arrowheads), shortest for falling phases (90 deg, marked with orange arrowhead), and increase progressively between 90 deg and 360 deg. Medians across rats are shown in black with 95% bootstrap confidence intervals on the within-rat effect of reference phase. C) Subtracting the mean delay for each rat from the lines shown in B to focus on the within-rat effect of reference phase on delay shows a reliable positive correlation between reference phase and conduction delay. The black line shown the best-fit trendline (R^2^ = 0.82) corresponding to the shown equation. Across panels, data from individual rats shown as colored lines. Ref. = reference.

Because waveform asymmetries, even for an otherwise synchronized rhythm, would be sufficient to generate apparent phase offsets, we next analyzed how waveform changes along the dorsal-ventral axis may have contributed to the total phase shift along the long axis of MEC. To do so, the asymmetry index values were converted into phase offsets, indicating the expected phase difference for each electrode pair as a result of the differences in asymmetry (Figure 6B). Comparing the phase shifts expected from changes in asymmetry to the empirically observed phase shifts (Replotted in Figure 6A for ease of reference) revealed that the asymmetry-related-phase-shifts accounted for a substantial portion of the total observed phase shift. After accounting for the apparent phase offsets due to waveform shape, the residual phase shifts were relatively flat with respect to position along the dorsal-ventral axis (Figure 6C). Quantitatively, the remaining delays were significantly reduced (i.e., traveled with a faster velocity) from the empirical delays (Kruskal–Wallis *H* test, *χ* ^2^(2) = 9.579, *P* = 0.0083; Figure 6D). These data demonstrate that most of the total phase shift observed along the dorsal-ventral axis of the MEC can be attributed to theta waveform changes.

**Figure 6.**
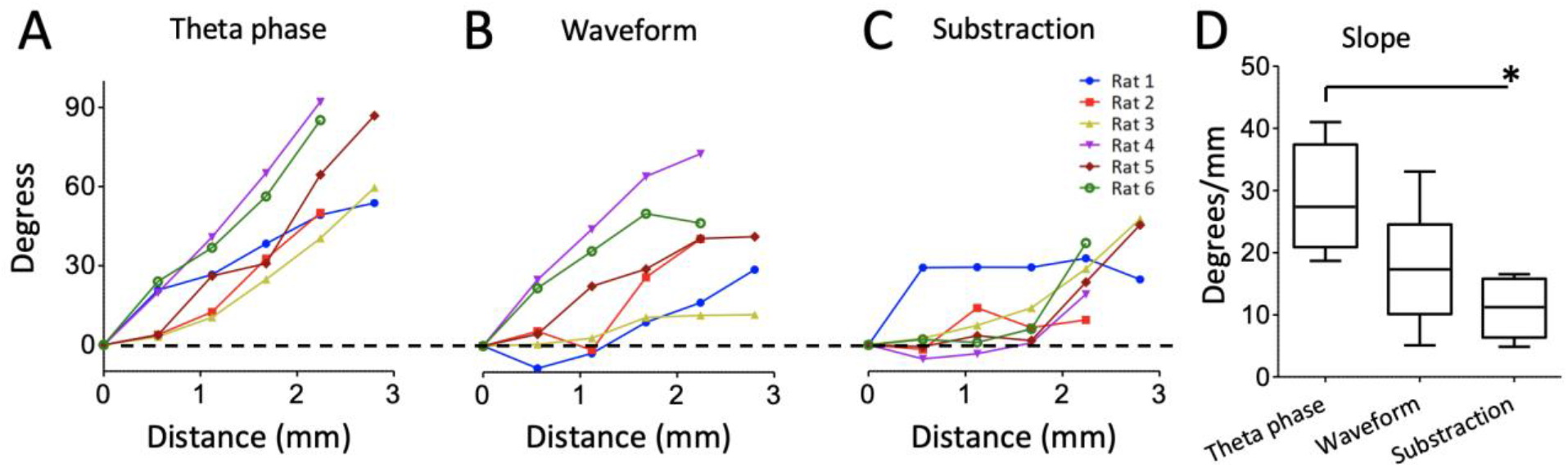
Theta waveform changes are a source of phase shift along the long axis of the MEC. A) Observed phase shifts across channels (i.e., theta-phase-shifts) replotted from Figure 2D for ease of comparison. B) Phase shifts expected given the changes in waveform asymmetry between sites (i.e., waveform-related-shifts). C) Subtraction of waveform-related-shifts from theta-phase-shifts across electrode positions shows remaining phase shifts are relatively flat with respect to distance along the dorsal-ventral axis. D) Comparison of the average phase shifts (degrees/mm) shown in A-C reveals a significantly reduced phase shifts after subtracting waveform-related-shifts from the theta-phase-shifts.

## Discussion

The goal of the work described here was to establish whether traveling waves in the entorhinal-hippocampal circuit are restricted to the hippocampus proper or whether the medial entorhinal cortex exhibits matched traveling waves. We answered this question by recording at regular intervals along the dorsal-ventral axis of the medial entorhinal cortex (MEC) in freely behaving rats. These recordings showed reliable phase shifts that were matched to those described previously to exist along the dorsal-ventral (septal-temporal) axis of the hippocampus. Paralleling these phase shifts were gradual changes in the theta waveform. These observations advance what is known about the basic mechanisms of cortical information processing, the potential function of theta, and the physiological basis of theta in the MEC.

The present work not only adds the MEC to the growing list of circuits recognized to exhibit traveling waves when engaged (for a recent review see Muller et al., 2018). Importantly, it is also the first to demonstrate that interacting functionally-distinct circuits have matched traveling waves. This is a necessary condition for traveling waves serve as a basic mechanism for coordinating interactions between distinct circuits. As such, the present work provides empirical support for existing theories regarding the utility of traveling waves for allowing the inherently distributed nervous system to operate as a coherent whole (Ermentrout and Kleinfeld, 2001; Muller et al., 2018).

Though the hippocampus and entorhinal cortex are neighboring anatomically the traveling waves observed here are not volume conducted from the hippocampus for a couple of reasons. First, laminar profiles of theta in the MEC demonstrate that theta is generated locally (Mitchell and Ranck, 1980). Second, the 90-degree phase shifts observed here are larger than would be expected from the limited extent of the hippocampus that neighbors the MEC. Rather, both areas have locally generated current sources that exhibit traveling waves.

The existence of matched traveling waves in the MEC and hippocampus suggest the possibility of a wide-spread network of brain areas for which graded phase offsets regulate the functional dynamics. The MEC and hippocampus are only two of a broad set of brain areas that exhibit theta rhythmic activity (Buzsáki, 2002). Indeed, prior models of theta function have presupposed such a distributed set of graded phase offsets and have shown that such offsets can afford the related network with adaptive functional properties related to navigation and memory (e.g., Blair et al., 2007; Burgess et al., 2007; Dickson et al., 2000; Hasselmo et al., 2009).

Crucial for linking function to physiology is the question of how this traveling wave is generated. We found that a substantial portion of the phase shifts observed across the MEC were related to differences in theta waveform. At dorsal sites, a sawtooth pattern was observed, characterized by a rapid rising phase and a gradual descending phase, whereas at ventral sites a more sinusoidal pattern was observed. Oscillation waveforms reflect the properties of their underlying physiological generators (for a review, see Cole and Voytek, 2017). The waveform changes we observed across the MEC indicates that the underlying generators vary along the dorsal-ventral axis. We hypothesize that the rapid rise of the sawtooth theta pattern observed at dorsal sites reflect an active local current source generated by the recruitment of local fast-spiking feedback inhibition (Pastoll et al., 2013). The same feedback inhibition may functionally support the formation of grid cell attractor networks (Pastoll et al., 2013). Notably, a prominent type of fast spiking feedback inhibitory neuron, parvalbumin expressing (PV+) interneurons, are distributed across the MEC in a gradient with high densities at dorsal sites and low densities at ventral sites (Beed et al., 2013). This distribution matches the anatomical distribution of the saw-tooth waveform pattern observed here and could account for the observed waveform changes. Alternatively, the waveform shifts could be related h-channels, which underlie theta generation (Kocsis and Li, 2004) in the medial septum and have graded strength along the dorsal-ventral axis of the MEC (Giocomo and Hasselmo, 2009). H-currents, too, are associated with the generation of grid tuning (Giocomo et al., 2011; 2007). Whether inhibitory tone or h-currents, that the same mechanism may underlie generation of the MEC traveling wave and grid cell attractor networks implies a deep functional connection between the traveling wave and spatial processing in the MEC.

In summary, our data demonstrate the existence of theta traveling waves in the medial entorhinal cortex that are matched to the traveling waves known to exist in hippocampus proper. The existence of matched traveling waves in interacting cortical areas supports the hypothesis that traveling waves are a basic organizing mechanism for coordinating cortical information processing. With regard to underlying mechanism, our findings that theta waveform varied in a graded fashion across the axis, indicate that the traveling wave is not generated by a propagating wave front but rather varying physiological generators across the axis, constraining mechanistic models of theta generation and function in the entorhinal-hippocampal circuit.

## Methods

### EXPERIMENTAL MODEL AND SUBJECT DETAILS

#### Animals

Recordings were made in 6 male Long-Evans rats. All animals weighed 350–500 g at the time of surgery, were individually housed, were maintained at 90% of their free-feeding weight following their full recovery after surgery. Animals were maintained on a 12:12 hr. light-dark cycls, all procedures were conducted during the light cycle.

### METHOD DETAILS

All experiments were performed in accordance with the National Institute of Health guide for the care and use of laboratory animals (NIH Publications No. 80-23) and were approved by the Bloomington Institutional Animal Care and Use Committee.

#### Electrode array construction

Custom multi-shank electrode arrays were built to record at regular intervals along the dorsal-ventral axis of the MEC. The positions of individual electrodes were fixed with respect to all other electrodes in the array. To build the electro array, 15-16 silica tubes (150 μm in diameter) were arranged and glued together in the horizontal plane. Individual shanks were created by loading stereotrodes or tetrodes (polytrodes) into single silica tubes or by merging the polytrodes loaded into adjacent silica tubes. Polytrodes were constructed by twisting two or four 12 or 25 μm diameter nichrome wires (A-M Systems) together. Five arrays were built to have two polytrodes per shank so as to have a vertical offset between the recording tips of the polytrodes within a shank of ∼280 μm. This design permitted us to use the deeper of the two as a scout electrode, seeking out the phase reversal that exists between layers II and I (Mitchell and Ranck, 1980). With this tip in layer I, the relatively shallow electrode was reliably positioned in layers II-III. The probe with up to six stereotrodes per shank, a vertical spacing of 450 μm was used to sample across the layers of the MEC more extensively (ED-Figure 2-1B).

Separate shanks were created in every other silica tube, resulting in a 300 μm separation between adjacent shanks. The total length of each successive shank from the silica increased by 470 μm. These spacings were selected so that the electrodes would track the ∼57° orientation of the MEC relative to the horizontal plane (Figure 1B). The net distance between the tips of adjacent shanks was net ∼566 μm. The full array of eight shanks spanned ∼3.9 mm in the dorsal-ventral axis of MEC.

Precise vertical spacing of stereotrodes within and between shanks was accomplished through the use of a micrometer graduated to a scale of 10 μm mounted to a microslicer (Stoelting part # 51425). In one array where neighboring shanks were glued together to create half as many shanks, the net resulting spacing was measured through inspection of array under a dissection microscope equipped with a reticle (Amscope part # EP10X30R).

Assembled arrays were connected to an electrode interface board and anchored to a single hand built microdrive (Vandecasteele et al., 2012). Finally, electrode tips were gold plated with a golden solution diluted 90% with a solution of 1 mg/mL polyethelene glycol in distilled water to bring the impedance at 1 kHz down to ∼250 kΩ.

#### Electrode array implantation and positioning

For electrode array implantation, rats were anesthetized with isoflurane, and a local anesthetic (Lidocaine 2 mg/kg and Bupivicaine 1 mg/kg) was administered subcutaneously to the scalp before any surgical incision. After the scalp was gently retracted and the surface of the skull was exposed, 4 to 5 anchor screws and one ground screw (located over the central cerebellum) were affixed to the skull. A craniotomy was performed over right MEC, at 4.5 mm lateral and extending from the transverse sinus to ∼2.5 mm anterior of the sinus. The electrode array was aligned 0.3 mm anterior of the transverse sinus and lowered 4.5–5 mm into the brain at the time of surgery. The microdrive was attached to the screws and the skull with dental acrylic. A copper mesh was then used to fashion a protective cage around the drive and was solidified using dental acrylic (Vandecasteele et al., 2012).

The rats recovered for seven days after surgery before behavioral testing and recording began. The electrode array was then stepped down to the MEC by advancing the microdrive in 140 μm increments until theta activity was prominent over most electrodes of the array (typically 5.7-6.0 mm at the deepest electrode). The final placement was established when the ventral-most recording sites crossed the boundary between layers II and I indicated by a reversal of theta phase between the polytrodes of a single shank (Mitchell and Ranck, 1980).

#### Electrophysiological recordings

Recordings were performed using a 32-channel headstage (Intan Technologies part RHD2132) connected directly to a USB interface board (Intan Technologies part an RHD2000). Data acquisition was controlled via a variant of the OpenEphys software platform customized to allow sync-pulses to be triggered by the electrophysiology to be sent to the behavioral tracking camera system. Field potential recordings were referenced to ground and written to disk at 30 kHz sampling rate.

#### Behavioral protocol

All recordings were performed as rats completed laps on a circle track for pieces of sweet cereal in 25-min-long testing trials. The track was made from 8-cm-wide black acrylic with 5-cm-tall opaque black plastic walls on both the inside and outside edges (Figure 1A). The circle had a diameter of 112 cm and was elevated ∼ 100 cm from the floor by six legs. Animals were rewarded for each complete lap run in either direction. The rewards were delivered in a fixed position by an experimenter standing arm’s length from the track. Numerous landmarks (e.g., experimenter, desk, door, pedestal, shelving) were visible from the track in the recording room. Rats received at least two weeks of training, consisting of one or more 20 min trials each day, prior to the key recording trials.

#### Histology

At the end of the experiments, the rats were deeply anesthetized with isoflurane and intracardially perfused with buffered formaldehyde solution (4%). Then, the brains were removed from the skull and soaked in 30% sucrose solution until saturated. The brains were then sectioned in the sagittal plane into 40 μm slices. Slices were mounted to slides and stained with cresyl violet to identify electrode tracks. Electrode position at the time of recording was established by cross-referencing images of these slides to logs of electrode movements, allowing for corrections when electrodes were advanced further after the recording. Raw histological micrographs are shown as Figure S1.

### QUANTIFICATION AND STATISTICAL ANALYSIS

Analyses were performed in MATLAB using custom scripts and with the CMBHOME toolbox.

#### Electrode selection criteria

Which specific electrodes were used in the analyses was determined in a multistep selection process. First, any electrodes with unstable signal either because it was flat or because the impedance was abnormal (< 100 kΩ or > 8 kΩ at 1000 Hz) were excluded from consideration. Second, the position of the array relative to the layers of the MEC was estimated by cross-referencing the known probe geometry to the observed electrophysiological markers. With regard to electrophysiological markers, specific attention was paid to theta power, theta phase reversals and, when available, the phase locking of local spiking activity. Given these estimates, the polytrode positioned closest to layer III from each shank was selected. Finally, with these stereotrodes identified, one of the two electrodes from each polytrode as selected. The final selections were reviewed for accuracy with histological verification of the electrode pathway.

#### Data inclusion criteria

Analyses were performed on epochs of data that were free of artefacts and had prominent theta fluctuations unless otherwise noted. Artefacts were defined as voltage swings greater than five times larger than the root mean square (RMS) of the signal. Identified artefacts, along with a 500 ms buffer before and after the artefact, were marked to be excluded. Because artefacts inflate the RMS of the signal, potentially creating inappropriately high thresholds for recognizing artefacts, the artefact removal procedure was applied to the resulting ‘cleaned’ data repeatedly until no further artefacts were found. For electrodes with acceptable impedance, this usually resulted in < 5% of the data being removed. From the remaining data epochs with high amplitude theta were selected for further analysis. High amplitude theta was defined as when the amplitude of theta (extracted by a zero phase-lag 4^th^ order bandpass Butterworth filter with 6 and 12 Hz cutoff frequencies) was greater than 400 µV for 500 ms or longer. Any theta amplitude dips below this threshold that lasted for 100 ms or less were ignored.

#### Calculation of average theta waves

To establish a picture of the average alignment of theta across the dorsal-ventral axis, we computed event related averages of the raw LFP triggered on the peak of theta on the dorsal-most channel. Importantly, this procedure did not use filtered signals, thereby avoiding the possibility that variations in theta waveform were altered. The peak of theta on the dorsal-most channel was accomplished using the waveform-based estimation of theta phase (Belluscio et al., 2012; Cole and Voytek, 2017). Briefly, a broad bandpass-filter (1-25 Hz) was applied to the dorsal-most recording to detect local maxima and local minima. These were defined as the theta peaks and troughs, respectively.

#### Phase locking

To estimate the reliability of the phase offset between electrodes, we computed the mean resultant length of the phase differences between electrodes. The phase differences themselves were computed as the circular difference between the instantaneous theta phase estimates from each of the channels. Instantaneous phase estimates for this analysis were computed from Hilbert transformed bandpass filtered LFP (zero phase-lag 4^th^ order bandpass Butterworth filter with 6 and 12 Hz cutoff frequencies). This method was selected *a priori* over the waveform-based approach used elsewhere because the linear interpolation used by the waveform-based approaches to label phases between peaks and troughs introduces the risk of creating artificially consistent phase offsets between channels. Though the Hilbert-based method introduces the possibility of finding negative phase changes, allowing for this variability reflects a conservative approach in this context as we report that phase locking is extremely high.

#### Propagation velocity of theta peaks and troughs

The propagation velocity of theta peaks (or troughs) across dorsal-ventral axis of the MEC for each recording was defined as the median propagation velocity over all of the individual theta cycles of the recording. To perform this analysis, all theta peaks (or troughs) were found on each channel using the waveform-based method described above. For each peak (or trough) found on the dorsal-most channel we defined a cycle. The peak (or trough) from each of the other channels that had the smallest absolute temporal offset to the peak (or trough) on the dorsal-most channel was defined as coming from the corresponding cycle. Importantly, this did not introduce bias with respect to propagation direction. For each cycles, we then performed a robust regression (via Matlab function robustfit.m) between the estimated anatomical position of each channel along the dorsal-ventral axis and the temporal offset of each peak (or trough) relative to the reference peak on the dorsal-most channel. The resulting slope was taken as an estimate of the propagation velocity for that cycle. Cycles for which the regression failed to converge were dropped from the analysis. Between 3500 to 5000 individual theta cycles were analyzed in each rat.

#### Theta band coherence

Theta band coherence was calculated as the average coherence across the 6 to 12 Hz frequency band (1 Hz increments). The coherence values themselves were derived by pass the artefact-free data to the Matlab function cohere.m. Importantly, this analysis was not restricted to epochs of high theta amplitude.

#### Analysis of multi-unit spiking activity

To test if neuronal activity also exhibited traveling waves, the electrodes were pushed deeper to layer II as indicated by strong trough-locked multi-unit activity. Multi-unit activity was isolated from the broadband signal by bandpass filtering the raw LFP (zero phase-lag 4^th^ order Butterworth bandpass filter with 600 and 3000 Hz cutoff frequencies). Then to extract the theta-timescale modulations of this spiking band, we bandpass filtered the amplitude envelope of the resulting spiking-band activity to the 4-12 Hz band (zero phase-lag 3^rd^ order Butterworth filter). To test for systematic lags in the change of this theta modulated multi-unit activity, we computed the temporal cross-correlogram between each channel and the dorsal-most channel and, for each, extracted the lag to the maxima within a window of +/− 120 ms.

#### Asymmetry analysis

To test for changes in the shape of the theta waveform, we used the asymmetry index described by Cole and Voytek (2018). Briefly, theta peaks and trough were identified using the waveform-based method. The waveform asymmetry was evaluated based on the rise-decay symmetry bases, the ratio of the rising phase duration to the decaying (falling) phase duration. The log10 transform of this ratio was then taken to create the index. This asymmetry index was computed for each theta cycle. The median across all theta cycles was then used to reflect the waveform asymmetry for that channel for that recording.

To estimate the phase offset that would be expected given specific theta waveform asymmetries, the difference in the duration of the rising and falling phases for each theta cycle was divided by that cycle’s trough-to-trough period. The resulting quotient was then multiplied by 360°.

#### Comparison of propagation rates for theta power and theta phase

To evaluate the propagation rate of theta power and theta phase along long axis on MEC, we used an approach that examined the lags to the maximum peak in the corresponding cross-correlograms. This analysis required a different approach from that used to calculate the propagation velocity of the theta peaks and troughs (described above) because theta power fluctuations do not have sufficiently regular peaks and troughs to use the prior approach here. Likewise, the present approach would not have worked for calculating peak and trough propagation velocity as it considers the whole signal at once instead of as distinct phases.

To estimate the propagation rate for theta phase, we computed the cross-correlation between theta activity recorded on the dorsal-most channel and on each of the other channels. Theta activity in this analysis was derived via a bandpass filtering of the raw LFP (zero phase-lag 4^th^ order bandpass Butterworth filter with 6 and 12 Hz cutoff frequencies). For each of the resulting cross-correlograms, we found the temporal lag to the local maxima closest to 0. We then performed a robust regression relating the spacing between the channels to the observed lag. The slope derived from this regression was taken as the propagation rate for that animal. A similar approach was taken for calculating the propagation rate for theta power changes. The only difference was that rather than calculating the cross-correlation between channels with respect to theta, we calculated it with respect to the amplitude of theta.

#### Statistics

Nonparametric statistics were used throughout the manuscript. In the case of nonpaired tests, Wilcoxon rank-sum tests were performed, In the case of paired tests, Wilcoxon signed-rank tests were performed. A nonparametric one-way Kruskal-Wallis was used to compare the phase shift related to waveform, subtraction of waveform and total phase shift, and a Dunn’s multiple comparation was performance as post hoc test. Significance was defined by an alpha level of 0.05.

### DATA AND SOFTWARE AVAILABILITY

All data are available by request to the authors.

## KEY RESOURCES

| Software and Algorithms |  |  |
| --- | --- | --- |
| OpenEphys GUI |  | http://www.open-ephys.org/gui |
| MATLAB |  | https://www.mathworks.com/products/matlab.html |
| CMBHOME toolbox |  | https://github.com/hasselmonians/CMBHOME |

## Author Contributions

J.H. Performed research, analyzed data, and wrote the paper

K.C. Analyzed data

E.N. Designed research, analyzed data and wrote the paper

## Acknowledgements

We thank the Whitehall Foundation and CONACYT (232364) for their generous support of this project. We are also grateful to J. Hinman, M. Brandon, & S. McKenzie for their comments and suggestions. Finally, we thank the staff of Indiana University laboratory animal resources for their care and attention to the subjects of this work.

## Supplemental Information

**Figure S1.**
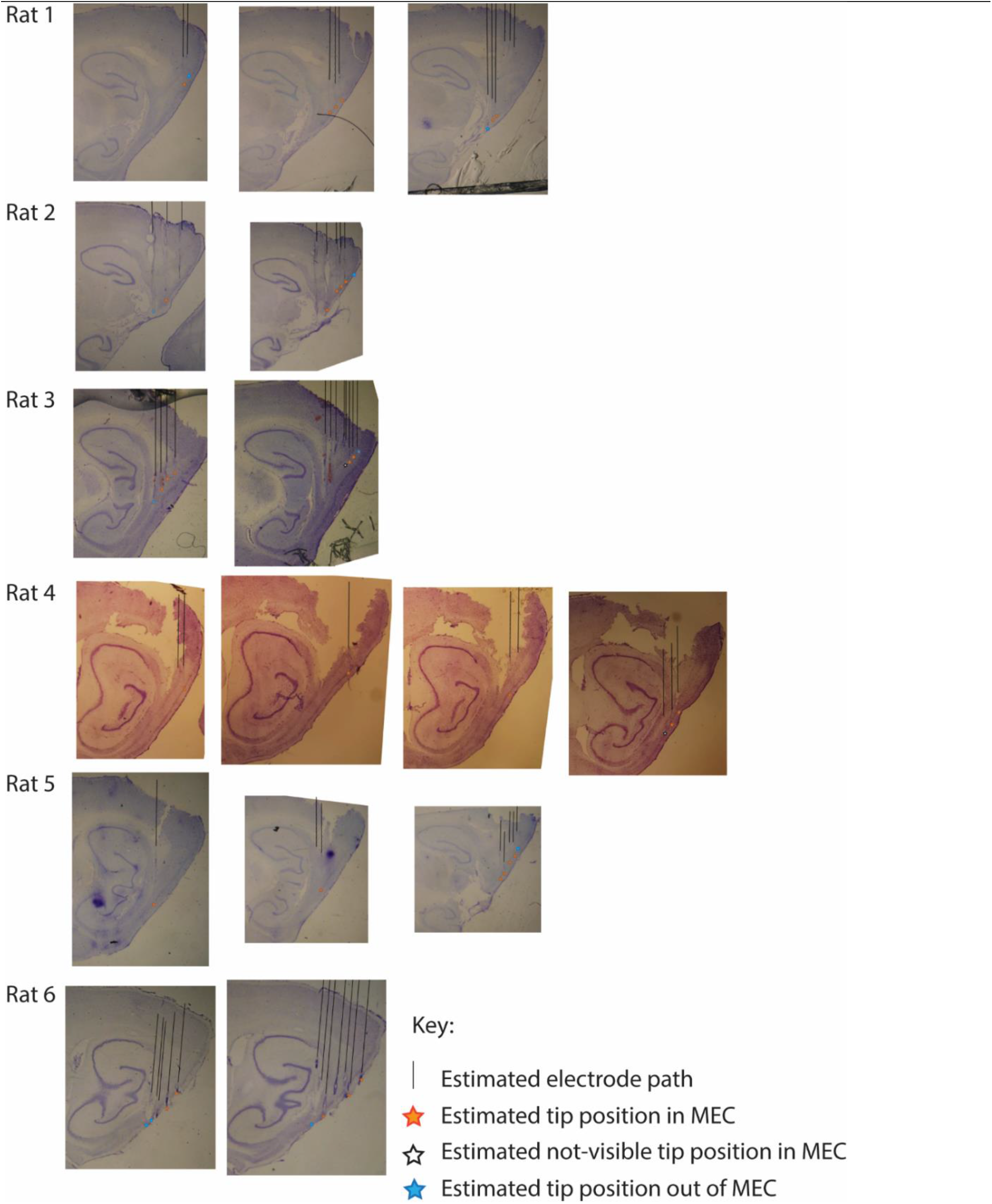
Histological micrographs for the six rats implanted for this study marking estimated final electrode placement at the time of sacrifice. Inferred positions at time of recording are shown as green stars. Black lines were added to reflect indicate the inferred electrode track. Stars were added to indicate the inferred final electrode position, blue reflects electrodes found to be outside of the medial entorhinal cortex, orange reflects electrodes found to target the medial entorhinal cortex. Note, in most cases, key recordings for which analyses upon which analyses were performed are relatively dorsal from the final termination sites shown here as electrodes were often stepped deeper in an effort to get subsequent recordings of unit activity in layer II.

**Figure S2.**
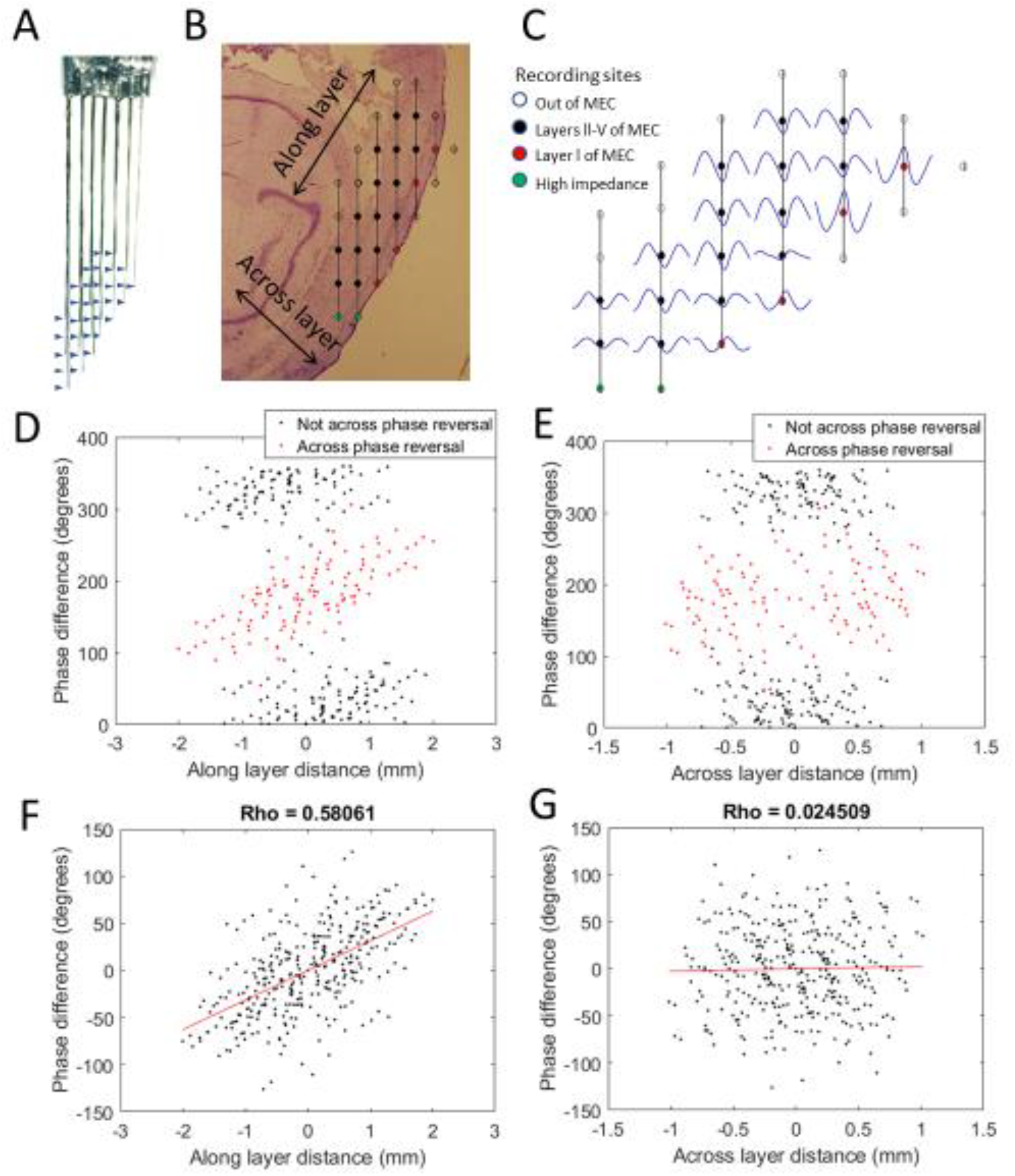
Progressive theta phase shift expressed along layer but not across layers in MEC. A) Custom multi-shank electrode array with tips positioned to allow simultaneous recording along and across the cortical layers of the MEC. Arrow heads indicate position of recording sites on adjacent shank. B) Reconstructed position of the electrode array based on histology, probe geometry, and electrophysiological markers. Color coding of individual sites indicates whether they were excluded for either being beyond the bounds of the MEC (white) or having high impedance (green) and whether they were found to have a phase reversal (red) relative to the remaining sites (black). C). Cycle-average theta waves for all electrodes, triggered off of peaks on the most dorsal electrode of layer l. The progressive theta phase shift is observed along the same layer, but not across layer, note the phase inversion of 180° between layer I vs layers III-V. D) The theta phase difference for each electrode-pair plotted as a function of the distance between the corresponding electrodes along the cortical layers (as shown in B) reveals a bimodal distribution that separately cleanly based upon whether the electrode pair spans the phase reversal expected between layers II and I (red) or not (black). E) Same as D but phase differences were plotted as a function of distance across the cortical layers (as shown in B). F) Applying a 180° offset to the red dots shown in D collapses the multi-modal distribution into a single modal distribution with a strong positive correlation between distance along the cortical layer and phase difference. G) Applying a 180° offset to the red dots shown in E also collapses the multi-modal distribution into a single modal distribution but with no notable correlation between distance across layers and phase difference.

